# Genetic Association of *LDHA* Variant g.2582481G>A with Racing Performance of Pakistani Pigeons

**DOI:** 10.1101/2022.08.14.503877

**Authors:** Rashid Saif, Muhammad Osama Zafar, Muhammad Hassan Raza

## Abstract

Pigeons are being used as a means of communication back in history due to their efficient flight stamina, navigational (homing) ability and endurance to harsh weather conditions. Due to these distinctive characteristics, pigeon racing and rearing has become a popular sport and hobby throughout the world, predominantly in Pakistan. The genetic tendency of *LDHA* gene in pigeons should be considered for understanding racing ability. The present study aimed to examine the genetic association of *LDHA* variant g.2582481G>A with racing performance in Pakistani pigeons. Our results demonstrated that 55% of the pigeon’s population was homozygous wild-type (G/G), 37.5% was heterozygous (G/A) and the remaining 7.5% was homozygous-mutant (A/A) at this locus. Furthermore, alternative allele frequencies within the best and low-performing cohorts of pigeons were 0.45 and 0.075 respectively. Furthermore, the Chi-Square association test was applied using PLINK data analysis toolset which showed a significant *p*-value of 1.381×10^-4^ with odds ratio (OR) of 10.09. Consequently, Pakistani pigeon population suggests a significant association of g.2582481G>A with racing performance similar to the other pigeon population of the world.

## Introduction

*Columba livia* has been in close interaction with humans due to its various aesthetic traits. Its domestication thought to be originated from Middle East and North Africa. Almost 300 different species of pigeon have been identified, out of which most popular pigeons are those who possess homing memory and spatial navigational traits (1). *Columba livia domestica* (homing pigeon) is derived from Rock pigeon due to some of its other traits as well e.g., docility, meat and droppings usage as fertilizer.

Pigeon has been used as rider in war back in the history due to its racing capabilities. Similarly, in the current era, masses are interested in arranging different tournaments in Pakistan and regional countries, making this bird more valuable for pigeon fanciers (2). In spite all of its morphological traits of keratinization and span of the wings, height and edging of keel bone, genetic factors are yet to explore in the selection of best racing pigeon. Several genetic studies reported the flight performance of pigeons can be linked to either environmental, weather, training method, nutrition and genetic factors (3).

Several genes e.g., *LDHA, DRD4, F-KER* and *MSTN* (4–6) markers have been identified and associated with racing performance of pigeons. Among the aforementioned, Lactate dehydrogenase-A (*LDHA*) gene is the most associated which encodes lactate hydrogenase-A enzyme. *LDHA* express in skeletal muscles and involve in the conversion of lactate into pyruvate (7). During long flight of pigeons, skeletal muscles undergo anaerobic respiration due to more energy requirement and limited supply of the oxygen. This accelerates the production of lactate in muscle fibers. Accumulation of the excessive lactate causing muscle fatigue and consequently decrease in pigeon speed and flight performance. LDHA enzyme regulates anaerobic respiration by converting lactate into pyruvate with reduction of NAD^+^ to NADH which may enhance the flight performance of pigeons (8).

The accession ID of *LDHA* gene in *Columba livia* is NW_004973198.1 with assembly of Cliv_1.0 (GCF_000337935.1). It consists of 8 exons and 7 introns having total length of 6,578 nucleotides, which encodes 332 amino acids (2). From the previous genetic studies, three SNPs had been reported having association with pigeon flight performance. Out of which, the 7^th^ intronic region of *LDHA* contains the highly associated marker g.2582481G>A which is significantly linked with pigeon’s flight performance. The mutant allele of this variant enhances the enzymatic activity of *LDHA* causing actively conversion of lactate into pyruvate (9). Without its accumulation in muscles, there are less chances of fatigue causing the increase in pigeon flight and stamina. Furthermore, the wild-type (GG) genotype is associated with low performance whereas heterozygous (GA) genotype give better performance in short distance races and homozygous mutant type (AA) are considered to be top performers.

The present study aims to detect the association of *LDHA* variant g.2582481G>A with pigeon flight performance by analyzing genotypic and allelic frequencies in the best and low performing pigeon cohorts.

## Materials and Methods

### Sample Collection and DNA extraction

To investigate genetic association of *LDHA* variant (g.2582481G>A) with racing performance of pigeon two cohort was selected for sampling from private pigeon fanciers who participated in this research project voluntarily. One cohort was assigned as low performing pigeons while other with best flight performance. Samples from each of the pigeon was taken independently from separate race to study genetic variability. Total 40 samples were taken with 20 from each cohort. Feather samples were taken and GDSBio (http://www.gdsbio.com) genomic DNA extraction kit was used to extract DNA from feather calamus.

### Primer designing for ARMS-PCR

Primers were designed to amplify *LDHA* wild and mutant alleles along with internal control against the transcript ID XM_005499258.2. OligoCalc was used to design and compute parameters of primers. For this purpose, total of 5 primers were designed, which were categorized as forward common, reverse normal (N) ARMS, reverse mutant (M) ARMS and two internal controls (IC). To increase specificity for *LDHA* genotyping a secondary mismatch was introduced at −3 nucleotide position from 3’ end of reverse ARMS primer. The details of primers are displayed in table 1.

**Table 1:**
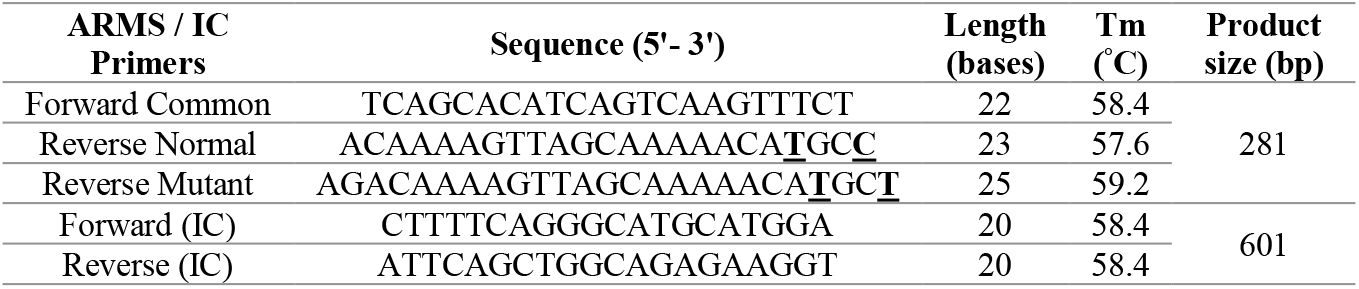
Details of primers sequences

### DNA amplification

ARMS-PCR reaction was performed using SimpliAmp thermal cycler (Applied Biosystems). Two separate reactions were made each labelled as Normal (N) and Mutant (M). Reaction for mutant was carried out by adding reverse mutant ARMS primer with common forward along with primers for internal controls. While for wild, reaction was made with normal reverse primer. A total of 25μL reaction mixture was prepared which contains 1μL genomic DNA, 10 mM of each primer, 0.05 IU/μL *Taq* polymerase, 2.5mM MgCl2, 2.5mM dNTPs, 1X Taq buffer, and PCR grade water. The PCR protocol was conducted with initial denaturation at 95°C for 5 min, followed by 30 cycles of denaturation (95°C for 30s), annealing (60°C for 30s) and extension (72°C for 30s). After that final extension was at 72°C for 10min.

### Statistical analysis

Chi-square association test with PLINK data analysis toolset was used to calculate the genotypic percentages, alternative allele frequencies, Odds Ratios and *p*-values of the subject variant.

## Results

### Variant genotyping

A total of 40 pigeons were genotyped. Out of which, the 20 pigeons were of best performing cohort while the remaining 20 were from low performing cohort according to the pigeon fanciers record. All these samples were collected from the local pigeoneers who trained them and took part in racing competitions at local level. The ARMS-PCR results showed significant association of g.2582481G>A with racing performance of our sampled pigeons. From the best cohort, 5 pigeons are G/G homozygous wild genotype, 12 pigeons are G/A heterozygous and 3 pigeons are A/A homozygous mutant. While from low cohort, 17 pigeons are G/G wild-type, 3 pigeons are G/A heterozygous with no homozygous mutant. The prevalence of alternative allele (A) was observed more in best performing cohort rather than low performing which indicates the association of g.2582481G>A with racing performance. The heterozygous and homozygous mutant genotypes pigeons may enhance the tendency of *LDHA* to convert the lactate into pyruvate that leads them to be the best racing performers. Figure 1 shows the ARMS-PCR amplification results of *LDHA* variant along with sampled pigeons’ pictures of the best cohort.

**Figure 1:**
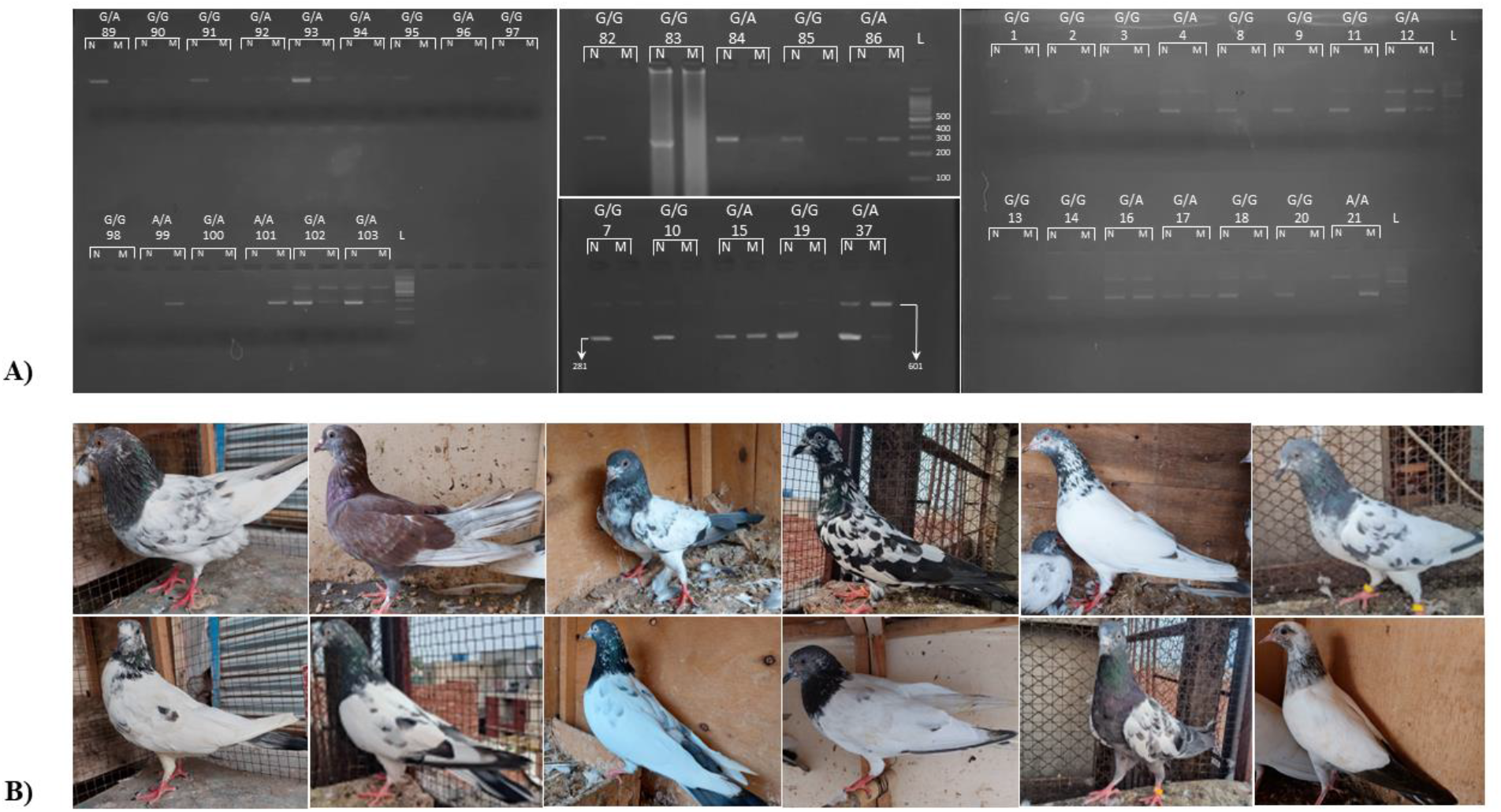
**A)** ARMS-PCR amplification of targeted regions N for wild-type and M for mutant-type samples **(B**) Sampled pigeons from the best performing cohort

### Association analysis

The genotypic percentages, allelic frequencies and *χ*^2^ association were calculated using PLINK data analysis toolset. 55% of the population are homozygous wild-type (GG), 37.5% are heterozygous (GA) while the remaining 7.5% are homozygous mutant (AA). Moreover, the alternative allele frequency (A) in best performing cohort is 0.45 comparatively higher than that of low performers 0.075. Chi-square significant *p*-value and odds ratio is 1.381×10^-4^ and 10.09 respectively. The association analysis of *LDHA* variant in Pakistani pigeon population is shown in table 2.

**Table 2:** Association analysis of *LDHA* locus in Pakistani pigeon population

| Scaffold / Position | cDNA Variant<br>XM_00549925<br>8.2 | Protein variant<br>XP_005499315.1 | Genotypic Information | | | Alternative Allele<br>Frequency | | $p$-Value<br>&<br>(OR) |
| --- | --- | --- | --- | --- | --- | --- | --- | --- |
|  |  |  | Wild-type<br>GG/% | Hetero<br>GA/% | Mutant-type<br>AA/% | Best<br>Performer | Controls |  |
| NW_004973198.1/<br>g.2582481 | c.999G>A | P.=<br>(Stop codon) | 22/55 | 15/37.5 | 3/7.5 | 0.45 | 0.075 | $1.381 \times 10^{-4}$<br>(10.09) |

## Discussion

*LDHA* gene variant g.2582481G>A showed substantial association with pigeon racing performance in *Columba livia* and considered as the highest contributing factor in worldwide pigeons population (7, 9). In the current research, we investigated the genetic association of the same variant in Pakistani pigeon population which showed the significant association in conformity to the global pigeon population. However, the sample size is small as compared to the other equivalent studies (8).

Our results presented that the overall twenty-two pigeons have homozygous G/G genotype, and the occurrence of G allele is relatively high while homozygous mutant (A/A) genotype was very rare in our sampled pigeon population. A/A genotype upregulates the expression of *LDHA* which increases the production of lactate dehydrogenase enzyme activity providing more energy to the bird’s wings muscle fibers to endure for the long time on flight (10). Consequently, lack of muscle fatigue and more energy consumption in muscle fibers leads to better racing performance.

Our genotypic association is linked to different research studies which are considered to determine the racing performance among the different pigeons breeds in Asia, Europe and Africa. A Japanese study on 123 pigeons was conducted in which three different SNPs of *LDHA* marker were screened i.e., g.2582481G>A, g.2583935G>A and g.2584057C>T and they concluded that particular one SNP, g.2582481G>A gave the highest association with pigeon performance in all type of races with significant *p*-value of 0.0107 (8). Another *LDHA* association studies from Poland showed genotypic (GG, GA, and AA) frequencies in different racing pigeons and their results demonstrated that AA genotype was more prevalent in top racers and this genotype was linked with racing performance having *p*-value of 0.001 (10). Similar analyses were conducted by Taiwan and Chinese researchers on 221 pigeons for the investigation of six polymorphic loci in *LDHA* gene which showed difference in genotypic distribution between the groups of homing and non-homing pigeons (7).

Genetic markers are not the only factor contributing to flight endurance but also, proper nutrition and training may affect their phenotypic traits. In future, other candidate genes as well as additional SNPs of *LDHA* needs to be scrutinize for complete picture and prediction for flying performance with large sample size.

## Conclusion

*LDHA* gene marker g.2582481G>A significantly associated with *p*-value of 1.381×10^-4^ in Pakistani pigeon population that can be used to predict the racing performance. This marker can be propagated further to make the subject trait more prevalent in the pigeon population by adopting the targeted marker assisted strategies to obtain desired population of the best performing pigeons to strive in harsh climatic condition in Pakistan.

## Ethical Statement

The ethical statement and approval were exempted for this study. No specific authorization from an animal ethics committee was required. Sampled pigeons belonged to private pigeon fanciers, who joined the present study on a voluntary basis.

## Acknowledgements

The authors acknowledge the pigeon fanciers and owners for sampling and joining this study.

